# Restricted effects of the sole *C. elegans* Daughterless/E homolog, HLH-2, on nervous system development

**DOI:** 10.1101/2022.10.10.511552

**Authors:** Neda Masoudi, Ralf Schnabel, Eviatar Yemini, Eduardo Leyva-Díaz, Oliver Hobert

## Abstract

Are there common mechanisms of neurogenesis used throughout an entire nervous system? Making use of the well-defined and relatively small size of the nervous system of the nematode *C. elegans*, we explored to what extent canonical proneural class I/II bHLH complexes are responsible for neurogenesis throughout the entire *C. elegans* nervous system. Distinct, lineage-specific proneural “class II” bHLH factors are generally thought to operate via interaction with a common, “class I” bHLH subunit, encoded by Daugtherless in flies, the E (E2A, E2-2, HEB) proteins in vertebrates, and *hlh-2* in *C. elegans*. To eliminate function of all proneuronal class I/II bHLH complexes, we therefore genetically removed maternal and zygotic *hlh-2* gene activity. We observed broad effects on neurogenesis, but still detected normal neurogenesis in many distinct neuron-producing lineages of the central and peripheral nervous system. Moreover, we find that *hlh-2* selectively affects some aspects of neuron differentiation while leaving others unaffected. While our studies confirm the function of proneuronal class I/II bHLH complexes in many different lineages throughout a nervous system, we conclude that their function is not universal, but rather restricted by lineage, cell type and components of differentiation programs affected.

## INTRODUCTION

Proneural basic helix-loop-helix (bHLH) transcription factors are phylogenetically conserved drivers of neurogenesis. Mutant analysis in flies and worms, as well as gain of function experiments in vertebrates revealed that members of this family are both required and sufficient for initial steps of neurogenesis (reviewed in (Jan and Jan 1994; Hassan and Bellen 2000; Bertrand *et al.* 2002; Wang and baker 2015; Baker and brown 2018)). Proneural bHLH factors fall into two families: the Achaete Scute family, members of which include the mouse Mash genes and Drosophila AS-C genes, and the Atonal family, which includes fly Atonal and its vertebrate Math orthologs, as well as *Drosophila* and vertebrate Neurogenin and NeuroD proteins (Hassan and bellen 2000; Bertrand *et al.* 2002; Baker and Brown 2018). Achaete Scute and Atonal family members are considered “class II bHLH” proteins that all heterodimerize with a broadly expressed, common “class I bHLH” protein (Massari and Murre 2000)(**Fig.1A**). As expected from the phenotype of their class II interaction partners, class I proteins (like fly Daughterless or the vertebrate E proteins E2, E2-2 and HEB) also have proneural activity (Wang and Baker 2015; Baker and Brown 2018). While proneural genes have been implicated in neurogenesis in many parts of invertebrate and vertebrate nervous systems, the extent to which canonical proneural class I/II bHLH complexes control neurogenesis throughout an entire nervous system has not been determined.

**Fig.1:**
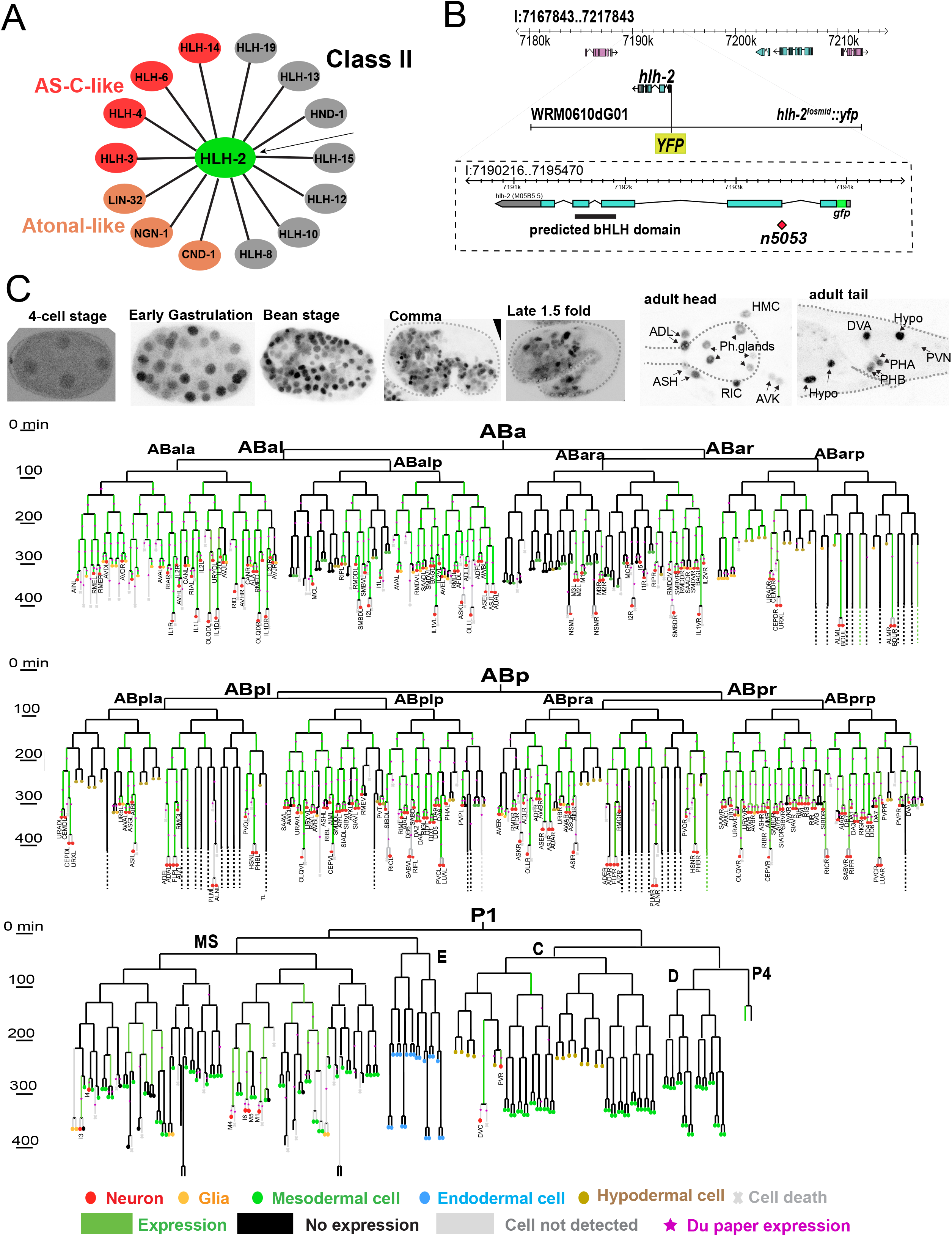
Background for HLH-2 protein function and description of its expression pattern. **(A)** Physical interaction of HLH-2 (Class I) with all Class II proteins as determined by Grove et al., 2009. Proneural AS-C and Atonal homologs are color-coded. No homodimerization of class II protein was detected. **(B)** Schematic of gene structure, showing mutation of G to A in the splice acceptor of second exon in n5053 allele and reporter genes. The YFP box shows the position of fluorescent reporter in the fosmid reporter (*otIs502*) and the gfp box represent the insertion of gfp, upstream of ATG in the CRISPR/Cas9-engineered reporter allele (*ot1089*), both of which show similar expression pattern. **(C)** Representative images of *hlh-2 (ot1089)* through out embryonic development (upper panel). The lower panels exhibit persistence of *hlh-2* signal in subset of neurons postembryonically. Dashed outlines indicate embryo shape. The lineage diagram indicates *hlh-2* fosmid expression (*otIs502*) in the AB and P lineages.

We sought to address this question in a comprehensive, nervous system-wide manner in the nematode *Caenorhabditis elegans.* The *C. elegans* genome codes for homologs of all genes classified as “proneural” in other systems (Bertrand *et al.* 2002)(**Fig.1A**). This includes three members of the Atonal family (a single Atonal ortholog *lin-32*, a single neurogenin ortholog, *ngn-1*, a single NeuroD ortholog, *cnd-1*), as well as five members of the AS-C family (*hlh-3, hlh-4, hlh-6, hlh-14, hlh-19*)(Ledent *et al.* 2002; Simionato *et al.* 2007)(**Fig.1A**). Proneural functions have been identified for several of these class II genes and, as expected, these proneural functions have been shown to also involve the sole *C. elegans* ortholog of the class I bHLH heterodimerization partner *hlh-2*/ Daughterless (Zhao and Emmons 1995; Krause *et al.* 1997; Portman and Emmons 2000; Frank *et al.* 2003; Nakano *et al.* 2010; Poole *et al.* 2011; Luo and Horvitz 2017; Masoudi *et al.* 2021), a notion further confirmed by biochemical interaction analysis (Grove *et al.* 2009)(**Fig.1A**). However, a comprehensive analysis of proneural genes in neuronal fate induction has been missing. The specific advantages that *C. elegans* brings to this problem is its nervous system-wide perspective: all *C. elegans* neurons are precisely mapped out, its neuron number is limited (302 neurons in the hermaphrodite) and molecular markers exist for each individual neuron class, thereby allowing to comprehensively probe proneural bHLH function.

Here, we provide a universal view of canonical proneural class I/II bHLH complex activities by analyzing the effects of the removal of the single class I bHLH ortholog, *hlh-2* (Krause et al. 1997). Based on the obligate heterodimer formation observed for all *C. elegans* proneural bHLH proteins (Grove *et al.* 2009), the genetic removal of *hlh-2* is expected to disable the function of all proneural bHLH genes. This would address the question to what extent proneural bHLH genes can be made responsible for the generation of all neuronal cell types in *C. elegans.* A similar approach has not yet been taken in other organisms. In *Drosophila* larvae, Daugtherless is required for the specification of many neurons of the peripheral nervous system (Caudy *et al.* 1988; Vaessin *et al.* 1994; Wang and Baker 2015). However, this conclusion is tempered by the limited number of molecular markers examined. Moreover, the maternal contribution of Daugtherless could not be removed without affecting reproduction, therefore leaving it unclear whether Daugtherless may have broader roles in the differentiation of the CNS of the fly. There are three class I genes in vertebrates, E12/E47 (aka TCF3), E2-2 (TCF4) and HEB (TCF12), but their function in neurogenesis, either in single or compound mutants has not been comprehensively analyzed to date (Wang and Baker 2015).

## RESULTS AND DISCUSSION

### Expression pattern of GFP-tagged HLH-2/Daughterless protein

Previous HLH-2 protein expression analysis, using antibody staining, showed transient expression in many parts of the developing embryo, as well as sustained expression in a few postembryonic cell types (Krause *et al.* 1997). However, the identity of many of the expressing cells remained unclear or tentative. We have used a chromosomally integrated fosmid reporter strain, *otIs502*, in which *hlh-2* is tagged at N-terminus with *yfp* (Sallee *et al.* 2017), as well as a CRISPR/Cas9 engineered reporter allele, with a N-terminal *gfp* insertion (*ot1089*) to precisely delineate *hlh-2* expressing cells (**Fig.1B**). For embryonic cell identification, we used 4D microscopy in conjunction with Simi BioCell software to trace *hlh-2* expression during embryonic development (Schnabel *et al.* 1997). Consistent with antibody staining (Krause *et al.* 1997), we detected low levels of HLH-2∷GFP at very early embryonic stages (the earliest stage we could detect is the 4 cell stage). Signals markedly increase at different time points in different lineages (**Fig.1C**). Expression was eventually established throughout most of the developing embryo, including all neuron producing lineages (**Fig.1C**). The only exception is the ABarapa lineage, which gives rise to five pharyngeal neurons (MCR, I1R, I2R, I5).

Expression of *hlh-2∷gfp* in the nervous system is mostly transient. It becomes undetectable in the vast majority of postmitotic neurons toward the end of embryogenesis. Postembryonic expression is only observed in four sensory neuron classes (ADL, ASH, PHA and PHB) and four interneuron classes (RIC, AVK, DVA, PVN)(**Fig.1C**). *hlh-2∷gfp* is expressed in all the P neuroblast at the L1 and very early L2 stage, but subsequently fades.

Together with proneural class II bHLH expression patterns that we reported earlier (Masoudi *et al.* 2018; Masoudi *et al.* 2021), a complete picture of proneural bHLH expression patterns emerges. As summarized in **Table S1**, *hlh-2* shows the broadest expression throughout the embryonic nervous system, followed by the also very broadly expressed AS-C homolog *hlh-3*. The other known proneural AS-C homolog *hlh-14* is the next most broadly expressed bHLH. ATO superfamily members are more narrowly expressed than *hlh-14*. Among the ATO superfamily members, *ngn-1* and *cnd-1* are more broadly expressed than *lin-32/Ato*.

### Lineage analysis of maternal/zygotic *hlh-2* mutant animals

Unlike most of the other proneural class II bHLH genes, strong loss-of-function mutants of the class I gene *hlh-2* display completely penetrant embryonic lethality (Nakano *et al.* 2010). Since homozygous *hlh-2* mutants derived from heterozygous mother appear to arrest development at different embryonic stages, we sought to remove potential maternal contributions using an unstable transgenic array that contains the wild-type *hlh-2* locus (see Material and Methods). We selected animals that had lost the array in the germline-producing lineage of their mothers. Such *hlh-2* maternal/zygotic mutants (henceforth called *hlh-2^m/z^* mutants) indeed display a stronger phenotype than zygotic *hlh-2* mutants (see Material and Methods). We set out to perform a cellular lineage analysis of *hlh-2^m/z^* animals using 4D lineaging with SIMI Biocell software (Schnabel *et al.* 1997). We focused on the AB lineage from which most of the nervous system derives (several of the few pharyngeal neurons derived from the MS lineage were previously also found to require proneural bHLH factors, including *hlh-*2)(Nakano *et al.* 2010; Luo and Horvitz 2017).

We found that 68 neurons are not born in *hlh-2^m/z^* mutants; instead, their mother or grandmother is transformed to hypodermal cell fate. Like conventional hypodermal cells, these “*de novo”* hypodermal cells stop dividing after 9^th^ division, migrate to the surface of embryo and do not have any speckled nuclei that are characteristic of neuronal nuclei. Additionally, we have found 9 cells normally destined to become a neuron, instead undergo programmed cell death (apoptosis) in *hlh-2^m/z^* embryos, exhibiting button-like condensed nuclei (**Fig.2, Table S2, S3**). Furthermore, we have found that the nuclei of the glia-like GLRL/R in the mutant look more like neuronal nuclei, hence we propose that these cells might have also switched their fate in *hlh-2* mutant. Other notable fate switches include the transformation of tail spike cell and some of the glial socket and sheath cells to hypodermal fate (**Table S2, S3**). We also observed an additional 23 hypodermal cells in place of cells normally destined to die by apoptosis. Among cells fated to go through apoptosis we have also observed a few instead exhibited neuronal features (speckled nuclei) and some that even went through an additional division (**Fig.2, Table S2, S3**).

**Fig.2:**
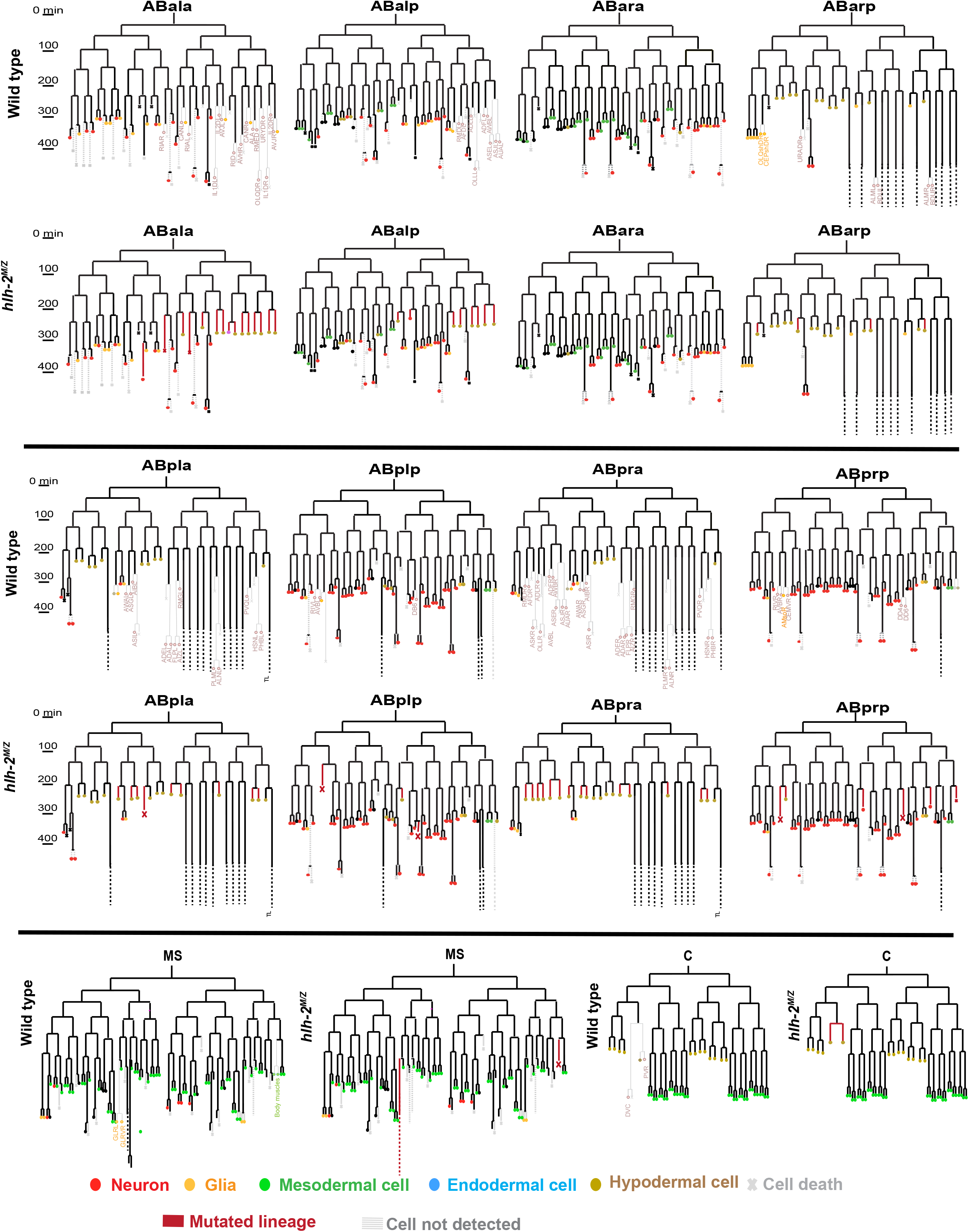
Lineage analysis of *hlh-2^m/z^* mutant animals. The break down of ABa, ABp, MS and C lineages are shown for detailed side-to-side comparison of wild type to *hlh-2(n5053)* mosaic progeny. The OH16873 strain was used for mosaic analysis. To eliminate any maternal contribution, hermaphrodites exhibiting the mosaic pattern that indicates loss of rescuing array in P2 lineage (*hlh-2^m/z^* mutant animals), were picked and their progeny were lineaged using SimiBiocell. Each lineage was analyzed thoroughly for timing and pattern of divisions, cell positions and potential cell fates based on morphological features using DIC imaging. The lineages affected are highlighted in the wild type panels with fader colors for ease of comparison. The presumptive fate is shown in the *hlh-2* panel, highlighted with red lines, right below every respective wild type section.

Taken together, our lineage analysis indicates that 122 of the 221 embryonically born cells fated to become neurons appear to be generated normally, as assessed by nuclear neuronal morphology, while 78 of the 221 embryonically born neurons are not generated and instead adopt alternative fates (either hypodermal or apoptosis) (**Fig.2, Table S2, S3**). **T**he remaining 21 neurons (221 in total, minus 122 neuron-like, minus 78 transformed) could not be lineaged or their morphological identity is ambiguous.

### Examination of hypodermal and neuronal fate markers in *hlh-2* mutants

We first set out to validate hypodermal transformations by examining the expression of a Zn finger transcription factor, *lin-26*, a pan-hypodermal cell fate marker with instructive roles in hypodermal cell fate specification (Labouesse *et al.* 1994; Labouesse *et al.* 1996). We generated a fosmid-based reporter for LIN-26 expression. Transgenic animals containing this reporter transgene (*otIs466)* reveal expression in ~50 more cells in *hlh-2^m/z^* mutant embryos, thereby confirming the generation of additional hypodermal cells (**Fig.3A**).

**Fig.3:**
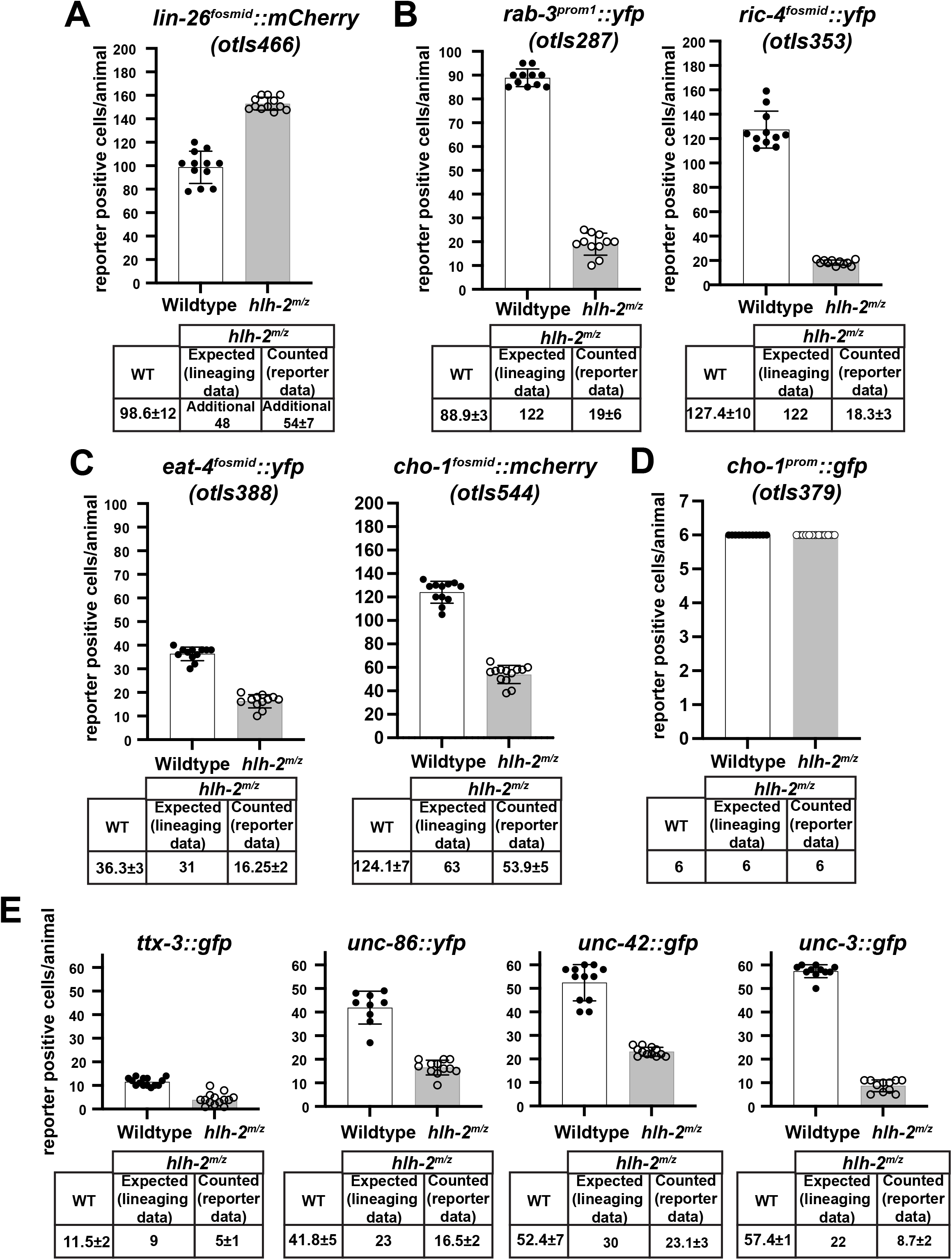
Effect of loss of *hlh-2* on cell type specific differentiation markers. **(A)** Effect of *hlh-2* removal on a hypodermal fate marker. **(B)** Effect of *hlh-2* removal on pan-neuronal reporter expression. **(C)** Effect of *hlh-2* removal on glutamatergic and cholinergic neuron differentiation. **(D)** Effect of *hlh-2* removal on a differentiation marker specifically expressed in AIA, AIY and AIN neurons. **(E)** Effect of *hlh-2* removal on a terminal selector markers.

Mirroring our analysis of pan-hypodermal fate, we examined the expression of two pan-neuronal marker genes, *rab-3* and *ric-4* (Stefanakis et al. 2015), in *hlh-2^m/z^* mutant embryos. We observed a striking reduction in the number of *rab-3∷yfp(+)* and *ric-4∷yfp(+)* cells, with less than 20 cells (of the 122 expected) expressing these two markers (**Fig.3B**). On the one hand, this observation confirms the lineaging observation that neurons are still generated in *hlh-2^m/z^* mutant embryos and therefore that proneural bHLH genes are not responsible for all embryonic neurogenesis. On the other hand, the extent of *rab-3∷yfp* and *ric-4∷yfp* marker expression loss (<20 cells still express these markers) is much more expansive than expected from the lineage analysis, which predicted that 122 neurons appear to be generated normally by light microscopical lineage analysis.

As an alternative means to visualize neuronal identity acquisition, we assessed the expression of neurotransmitter identity features. To this end, we examined expression of fosmid-based reporters that measure the expression of the vesicular transporter *eat-4*, a marker of glutamatergic neuron differentiation, and *cho-1*, a general marker of cholinergic differentiation (Serrano-Saiz *et al.* 2013; Pereira *et al.* 2015). These two neurotransmitter systems cover the vast majority of neurons in the embryonically generated nervous system and are expressed in neurons that our lineage analysis suggests is unaffected in *hlh-2^m/z^* mutant animals, as well as in neurons that display lineage transformations (**Table S3**). We observed a ~50% reduction in the total number of neurons expressing either *eat-4* or *cho-1* in *hlh-2^m/z^* mutant animals (**Fig.3C**). Since we still detect ~16 *eat-4(+)* and ~54 *cho-1(+)* neurons, we conclude that many neurons still execute at least a part of their differentiation program in *hlh-2^m/z^* mutant embryos. We determined the precise molecular identity a subset of these unaffected neurons by examining the expression of an enhancer fragment derived from the *cho-1* choline transporter (Zhang *et al.* 2014). This fragment is expressed in six neuron (AIA, AIY, AIN neuron pairs) and is unaffected in *hlh-2^m/z^* mutant embryos, corroborating that these neurons differentiate normally in the absence of *hlh-2* (**Fig.3D**).

The most striking aspect of our analysis of neurotransmitter identity markers is that the total number of *eat-4*-positive and *cho-1*-positive cells (70) far exceeds the number of cells that still express pan-neuronal markers (<20), indicating selective effects of *hlh-2* on neuron differentiation. We sought to further investigate this discrepancy by turning to another set of marker genes, as described in the next section.

### Neuron type-specific effects of *hlh-2* on terminal selector expression

To further assess neuronal cell fate acquisition in *hlh-2^m/z^* mutant animals, we turned to another set of molecular markers. Key drivers of neuronal identity are neuron type-specific terminal selector transcription factors (Hobert 2008; Hobert 2016). These factors exert their activity shortly after neuron birth, initiating the expression of neuron-type specific gene batteries, but not pan-neuronal identity features (Hobert 2016). Moreover, in a number of different lineages, terminal selectors have been found to be downstream of class II proneural bHLH genes that presumably work in conjunction with the class I heterodimerization partner HLH-2 (Masoudi *et al.* 2021). We analyzed the expression of four phylogenetically conserved terminal selectors, *ttx-3/LHX2*, *unc-86/BRN3, unc-42/PROP1* and *unc-3/EBF* which together mark the proper differentiation of about half of the 221 embryonically generated neurons (Finney and Ruvkun 1990; Pereira *et al.* 2015; Reilly *et al.* 2020; Berghoff *et al.* 2021).

The terminal selector *ttx-3/LHX2*, a LIM homeobox gene, is continuously expressed in the AIY, AIN, AIA, NSM and ASK neuron pairs (Reilly *et al.* 2020), as well as transiently in the SMDD neuron pair (Bertrand and Hobert 2009), a total of 12 neurons (**Fig.3E**). As per our light microscopy SIMI/Biocell analysis, all these neurons appear normal in *hlh-2^m/z^* mutant embryos with the exception of the single ASKR neuron where a hypodermal cell fate transformation is observed. We find that in *hlh-2^m/z^* mutant embryos, *ttx-3* expression is reduced by about half, down to six cells (**Fig.3E**). We reverse-lineaged some of the *ttx-3(+)* cells in *hlh-2^m/z^* mutant embryos and found that two of those remaining are the AIY interneuron pair (see Methods). We were surprised by the result because previous RNAi of *hlh-2* led to defects in *ttx-3* expression (Bertrand and Hobert 2009; Filippopoulou *et al.* 2021), but we note that it is not unprecedented that RNAi-induced defects could not be reproduced with mutant alleles (Schmitz *et al.* 2007; Jimeno-Martin *et al.* 2022). As an independent means to assess AIY differentiation and effects on *ttx-3* expression, we examined the expression of a direct target gene of *ttx-3*, the choline transporter *cho-1* (Stefanakis et al. 2015). As noted above, expression of a *cho-1* enhancer fragment that is expressed in the AIY, and also AIN and AIA interneurons, is unaffected in *hlh-2^m/z^* mutant animals (**Fig.3D**). We conclude from this analysis that in some neurons where there appears to be no lineage defects *hlh-2* indeed has no effect on differentiation (AIY, AIA, AIN), but that in other neurons (likely the remaining NSM, ASK, SMDD neurons), *hlh-2* affects neuronal differentiation based on the reduction of *ttx-3*-positive neurons.

A similar picture emerges when considering expression of the *unc-86* POU homeobox gene - another terminal selector that acts in multiple neuron types, in combination with other homeobox genes, to specify their identity (Leyva-Diaz *et al.* 2020). In the embryo, *unc-86* has been reported to be selectively expressed in 46 neurons (Finney and Ruvkun 1990), a number we confirmed with our reporter transgene (**Fig.3E**). Lineage analysis predicts a transformation of 23 of the neurons to hypodermal fate or programmed cell death, with the other 23 neurons appearing to be generated normally. We observe ~16 neurons to express *unc-86* in *hlh-2^m/z^* mutant embryos. As with *ttx-3*, this again indicates that many cells indeed differentiate appropriately to express *unc-86*, but because only 16 out of an expected 23 cells do so, we conclude that a subset of those neuron-like cells fail to differentiate properly (**Fig.3E**).

The *unc-42* homeobox gene is selectively expressed in 42 neurons during larval and adult stages (Berghoff *et al.* 2021) and our lineaging predicts 12 of these neurons to be transformed to hypodermal fate or programmed cell death (**Table S3**). Analysis of an *unc-42* reporter allele (*ot986)* reveals more expression in wild-type embryos (>50 cells), indicating previously undescribed, transient *unc-42* expression in additional cells. Almost half these cells lose *unc-42* expression in *hlh-2^m/z^* mutant embryos (**Fig.3E**). Since it is not clear whether this reduction is due to loss of expression in these transiently UNC-42-positive neurons, we reverse-lineaged *unc-42(+)* cells and found that some of these UNC-42-positive cells in *hlh-2^m/z^* mutant embryos include ASH, AVK, SIB and SMD (12 of the 23 UNC-42(+) neurons). These neurons do not show lineage transformation in our embryonic lineaging experiments and we therefore again conclude that several neuronal cell types acquire their neuron-specific identities in *hlh-2^m/z^* mutant embryos, while others do not. We further confirmed this notion by using another marker for terminal differentiation of AVK, the FMRF-amide encoding *flp-1* gene, a target of UNC-42 (Wightman *et al.* 2005). We find *flp-1∷gfp (bwIs2* transgene) expression to be unaffected in 15/15 examined *hlh-2^m/z^* mutants.

Lastly, we examined expression of *unc-3* the sole *C. elegans* ortholog of the Collier/Olf/Ebf (COE), which acts as terminal selector in several cholinergic motor neuron types (Kratsios *et al.* 2011; Pereira *et al.* 2015). As in the case of *unc-42*, we observed more widespread embryonic expression of *unc-3* in the embryo (~57 cells) as predicted by its postembryonic sites of expression (34, of which 22 are supposedly normally generated in *hlh-2^m/z^* mutants). However, we observe an extensive reduction of *unc-3* reporter allele expression, with only ~9 UNC-3-positive cells observed in *hlh-2^m/z^* mutant embryos (**Fig.3E; Table S3**). This again indicates that while many neuron-like cells fail to properly differentiate in *hlh-2^m/z^* mutant embryos, others do.

## CONCLUSIONS

Extensive work on neuronal cell fate specification in *C. elegans* has revealed some themes that are broadly applicable throughout the entire *C. elegans* nervous system. These include coordinated regulation of neuron-type specific terminal identity features by so-called terminal selectors, the separate regulation of neuron-type specific terminal identity features from pan-neuronal gene regulation, and the deployment of homeobox genes as neuronal identity specifiers throughout the entire nervous system of the worm (Hobert 2016; Hobert 2021). The existence of such universal themes prompted us to ask whether all *C. elegans* neurons also rely on a common mechanism for earlier steps of neuronal development, specifically, the decision to generate neuronal precursors from ectodermal-derived cells. In several different cellular contexts, so-called “proneural bHLH genes” (AS-C and Ath family members), were already known to fulfill such roles in many different animal species (Jan and Jan 1994; Hassan and Bellen 2000; Bertrand *et al.* 2002; Wang and Baker 2015; Baker and Brown 2018), yet a true nervous system wide perspective of the involvement of these proneural bHLH genes in neurogenesis was lacking. Through the removal of the common proneural complex subunit HLH-2, we have addressed here the question whether proneural bHLH genes can be made responsible for all neurogenesis in the *C. elegans* embryo. We draw the following three conclusions from our analysis (**Fig.4**):

**Fig.4:**
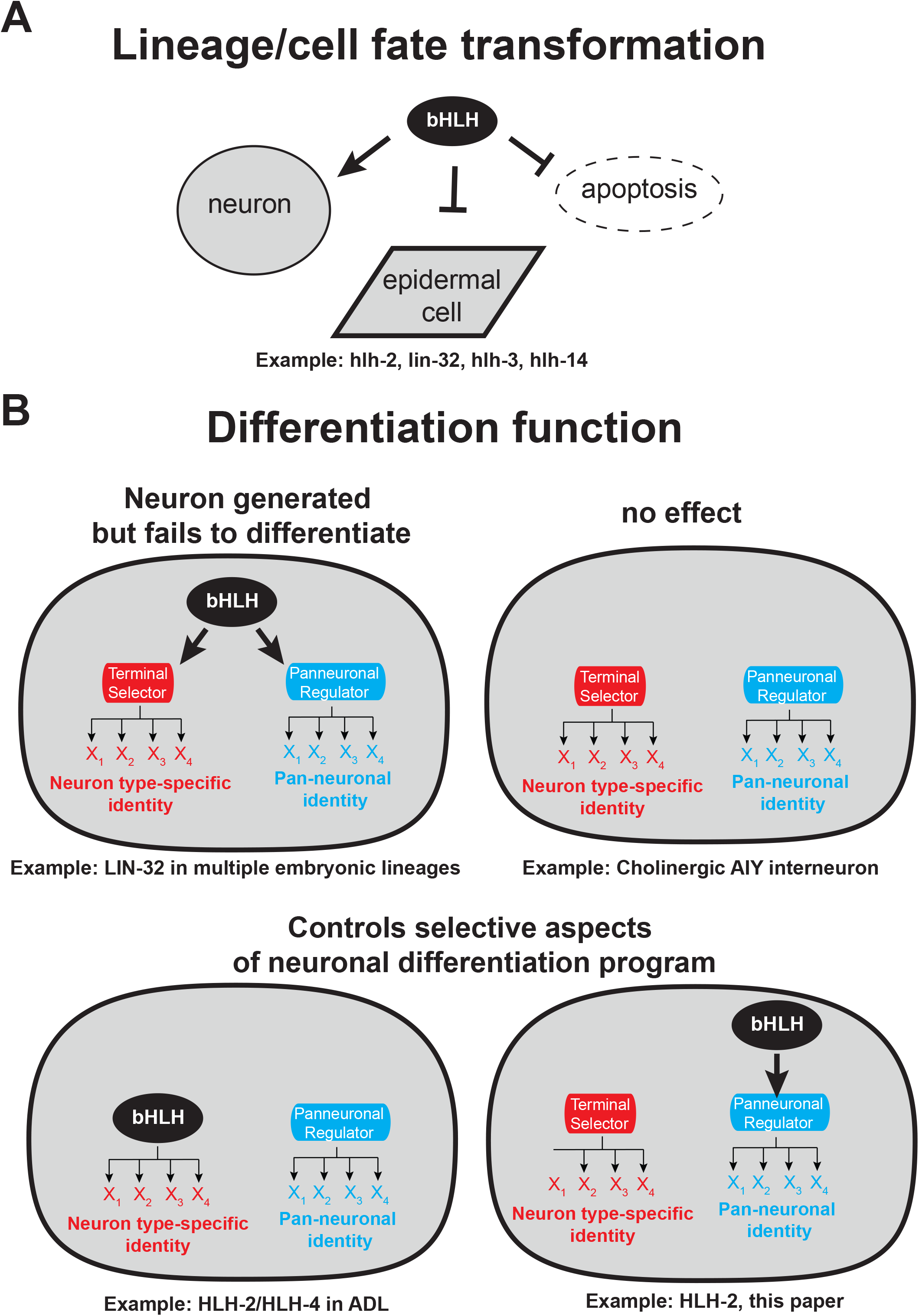
Summary of involvement of bHLH genes of the “proneural” bHLH family (class I Daughterless/hlh-2 and class II AS-C-like and Ato-like genes) in neuronal identity specification. **(A)** In class I (*hlh-2*) and several class II bHLH mutant backgrounds, “classic” proneural defects are observed, manifested by a transformation to hypo/epidermal cell fate (Zhao and Emmons 1995; Portman and Emmons 2000; Frank *et al.* 2003; Nakano *et al.* 2010; Poole *et al.* 2011; Luo and Horvitz 2017; Masoudi *et al.* 2021), analogous to proneural defects originally described in the *Drosophila* nervous system (Campuzano and Modolell 1992; Jarman *et al.* 1993). **(B)** Involvement of class I and class II “proneural” bHLH genes in neuronal differentiation in *C. elegans.* Neuronal differentiation is broken down here in genetically separable processes the control neuron-type specific gene batteries (via terminal selectors) and pan-neuronal gene batteries (via the CUT homeobox genes)(Leyva-diaz and Hobert 2022).

1. As expected, *hlh-2* – likely in combination with class II bHLH genes – does have a proneural function in many lineages throughout the central and peripheral nervous system of *C. elegans* with no particular preference for lineage, neuron position or neuron function. The most obvious manifestation of such role is the transformation of neuronal lineages or cells to hypodermal cell fates, a “classic” phenotype of proneural genes, initially observed in the peripheral nervous system of *Drosophila* (Jan and Jan 1994) and subsequently in *C. elegans* (Zhao and Emmons 1995). Such hypodermal transformations are only the most extreme version of a proneural function. Based on our previous analysis of *lin-32/Atonal* function (Masoudi *et al.* 2021), and confirmed here with our analysis of *hlh-2*, proneural functions are also manifested by a combined loss of terminal selectors, pan-neuronal markers, and neuron type-specific markers without concomitant switch to hypodermal identity.
2. Based on our lineage and marker gene analysis, we conclude that not all neurogenic processes in an animal nervous system require canonical class I/II proneural bHLH complexes. This is a novel conclusion that could not be made in any other system before and is due to our ability to comprehensively analyze neurogenesis throughout an entire nervous system. One may not have to look far to find alternative proneural factors. A recent report indicates that the combined removal of two conventional proneural AS-C/ATO-like genes (*hlh-3/AS-C* and *ngn-1/NGN*) in combination with an bHLH gene not previously considered to be a proneural factor, the OLIG-homolog *hlh-16*, affects the proper differentiation of the AIY interneuron class, as measured by expression of the terminal selector *ttx-3* (Filippopoulou *et al.* 2021). We found AIY to be unaffected by *hlh-2* removal. Both the redundant function of these factors (neither single mutant alone has a phenotype)(Filippopoulou *et al.* 2021), as well as their apparent independence of *hlh-2* suggests proneural function of non-canonical class II-only bHLH complexes.
3. There are striking discrepancies in the extent by which distinct types of neuronal marker genes are affected in *hlh-2* mutants, indicating that *hlh-2* has nuanced, cell-type functions in controlling select aspects of terminal neuron differentiation (**Fig.5**). Specifically, the effects of *hlh-2* removal on pan-neuronal genes appear to be broader than expected from the expression of neuron-type-specific identity markers (i.e. terminal selectors and other neuron type-specific effector genes). In other words, in several lineages, *hlh-2* may control pan-neuronal features independently of neuron-type specific differentiation programs. One limitation of our study is that we cannot pinpoint the nature of these neurons due to the overall morphological disorganization of *hlh-2^m/z^* mutant embryos. This conclusion therefore relies only on counting the number of fate marker-positive cells (e.g. pan-neuronal marker genes versus neuron type-specific marker genes) and comparing them to each other in wild-type and *hlh-2* mutant animals. In spite of this limitation, we note that this conclusion is consistent with previous reports that showed that pan-neuronal differentiation can be genetically separated from neuron-type specific differentiation. Specifically, terminal selectors affect neuron-type specific features, but leave pan-neuronal differentiation programs unaffected; conversely, members of the CUT homeobox gene family directly activate the expression of pan-neuronal identity features, but leave neuron specific identity features intact (Hobert 2016; Leyva-diaz and Hobert 2022). Hence, it is conceivable that HLH-2 complexes may either directly control pan-neuronal effector genes and/or may control the expression of CUT homeobox genes; however, this cannot be a universal function, since pan-neuronal features remain unaffected in several neuronal lineages.

In summary, nervous system-wide analysis of class I and class II bHLH gene function, reported here and in recent papers from our laboratory (Masoudi *et al.* 2018; Masoudi *et al.* 2021) provides novel and more nuanced views of proneural gene function in the nervous system of *C. elegans* that we summarize in **Figure 4**.

## MATERIAL AND METHODS

### Strains

Strains were maintained by standard methods (Brenner 1974). Previously described strains used in this study are as follow.

Mutant strains:

MT17677 - *hlh-2(n5053) / hT2[qIs48] I; + / hT2[qIs48] III* (Nakano *et al.* 2010)
MT19085 - *hlh-2(n5287) / hT2[qIs48] I; + / hT2[qIs48] III* (Nakano *et al.* 2010)
OH16873 - *hlh-2(n5053); otEx7684* (*hlh-2^fosmid^∷yfp; myo-3∷mcherry*)] (this paper)

CRISPR/Cas9-engineered reporter alleles:

OH16761 – *hlh-2[ot1089 (hlh-2∷gfp)* (this paper)
OH16111 - *unc-42[ot986 (unc-42∷gfp)* (Berghoff *et al.* 2021)
OH13990 - *unc-3[ot839(unc-3∷gfp)* (Kratsios et al. 2017)

Reporter transgenes:

*otIs502 - hlh-2^fosmid^∷YFP + myo-3p∷mCherry* (Sallee *et al.* 2017),
*otIs466 - lin-26^fosmid^∷mCherry; lin-44∷nls∷yfp* (this paper)
*otIs353 - ric-4^fosmid^∷yfp* (Stefanakis et al. 2015)
*otIs287 - rab-3^prom1^∷yfp* (Stefanakis et al. 2015)
*otIs388 - eat-4^fosmid^∷yfp∷H2B* (Serrano-saiz et al. 2013)
*otIs544 - cho-1^fosmid^∷SL2∷NLS∷mCherry* (Pereira et al. 2015)
*otIs379 - cho-1^prom^∷gfp* (Zhang et al. 2014)
*bwIs2 - flp-1∷gfp* (Much et al. 2000)
*otIs337 - unc-86^fosmid^∷NLS∷yfp* (Pereira et al. 2015)
*wgIs68 - ttx-3^fosmid^∷gfp* (Zhang et al. 2014)

### Maternal/zygotic *hlh-2* removal

The first indication that zygotic *hlh-2* mutants may not fully remove all gene activity was the variability in expressivity of arrest phenotypes that homozygous *hlh-2(n5053)* mutants, derived from heterozygous mothers displayed (Nakano *et al.* 2010). Animals arrested at different embryonic stages and some animals even survived till the first larval stage. To remove potential maternal contribution, we sought to rescue the embryonic lethality of homozygous *n5053* mutant animals with a transgenic array (*otEx7684)* that contains the fosmid carrying the entire *hlh-2* locus and its upstream genomic sequence. Due to their instability, animals can be identified via a co-injection marker expressed in body wall muscle (*myo-3∷mCherry)* that have lost the array in the P lineage (generating muscle and germline) of the parental generation and hence contain neither maternal nor zygotic gene activity. The effect of maternal contribution was most obvious when examining expression of the pan-neuronal *rab-3* marker. While zygotic mutants show no loss of *rab-3* expression (≈ 100), *n5053* homozygous animals derived from mother that did not contain the rescuing array in the germline forming P2 lineage (called *hlh-2^maternal/zygotic^* or *hlh-2^m/z^* mutants) showed a reduced number of *rab-3(+)* cells (≈20). demonstrating the impact of maternally supplied *hlh-2* on neurogenesis.

In spite of repeated attempts with multiple different genomic fosmid clones, we were not able to rescue the embryonic lethality of the *n5287* deletion allele. When crossed into *n5287* animals, even the transgenic array that rescued the *n5053* array could not rescue the *n5287* lethality. We can therefore not conclusively assign the embryonic lethality of *n5287* animals exclusively to the *hlh-2* locus and therefore used the *n5053* allele for all of our analysis.

### Microscopy

For fluorescence microscopy, worms were paralyzed by 25mM sodium azide (NaN_3_) and mounted on 5% agarose pad on glass slides. Images were acquired using an axioscope (Zeiss, AXIO Imager Z.2) or LSM 800 laser point scanning confocal microscope. Representative images are max-projection of Z-stacks. Image reconstruction was performed using Fiji software (Schindelin *et al.* 2012).

4D microscopy and SIMI BioCell was used, as previously described (Schnabel *et al.* 1997), to analyze embryonic lineage defects of mutant animals as well as bHLH fosmid reporter expression pattern during embryogenesis. Briefly, gravid adults were dissected on glass slide and a single two-cell stage embryo was mounted and recorded over 8 hours of embryonic development. Nomarski stacks were taken every 30 s and embryos were illuminated with LED fluorescence light (470 nm) at set time points during development. The recording was done with Zeiss Imager Z1 compound microscope, using the 4D microscopy software Steuerprogram (Caenotec).

## ACKNOWLEDGEMENTS

We thank Chi Chen for assistance with microinjections to generate strains, Ines Carrera for help with generating the *hlh-2* fosmid reporter, Luisa Cochella for comments on the manuscript and members of the Hobert lab for discussion. This work was funded by the Howard Hughes Medical Institute.

## SUPPLEMENTARY LEGENDS

**Table S1: Tabular comparison of all proneural bHLH expression profile**

The complete developmental expression patterns of the sole proneural Atonal ortholog *lin-32*, the sole NeuroD ortholog *cnd-1*, the sole neurogenin ortholog *ngn-1*, and three of the five AS-C homologs (*hlh-3, hlh-4, hlh-14*) has been generated using fosmid-based reporter transgenes and/or CRISPR/Cas9-engineered reporter alleles (Masoudi et al. 2018; Masoudi et al. 2021). The two remaining AS-C homologs, *hlh-6* and *hlh-19*, are not expressed in the nervous system. A 5’ promoter fusion reporter upstream of the hlh-6 locus is exclusively expressed in pharyngeal gland cells (Smit et al. 2008). To exclude the possibility that regulatory elements were missed in this reporter, we generated a fosmid-based reporter of *hlh-6* and also found it to be exclusively expressed in pharyngeal gland cells. We also generated a fosmid-based reporter for the previously unstudied *hlh-19* gene and found it to be weakly expressed in embryonic hypodermal cells but non in the developing nervous system. Tagging the endogenous *hlh-19* locus with *gfp* using CRISPR/Cas9, revealed no expression in any tissue.

**Table S2: Tabular summary of cell fate transformation to cell death or hypodermal fate in *hlh-2* mutants**. Transformations are grouped based on transformation to/ from cell death and to hypodermal cells. Also see **Table S3** for a summary of cell fate transformation, organized by neuronal cell type.

**Table S3: Tabular summary of neuronal fate transformation in *hlh-2* mutant.**

Transformations are highlighted with different color codes (Data is shown in Figure 2). The expression of all markers used in this study is also specified by color codes (Data is shown in Figure 3).

